# Separate *Polycomb Response Elements* control chromatin state and activation of the *vestigial* gene

**DOI:** 10.1101/488478

**Authors:** Kami Ahmad, Amy Spens

## Abstract

Patterned expression of many developmental genes is specified by transcription factor gene expression, but is thought to be refined by chromatin-mediated repression. Regulatory DNA sequences called *Polycomb Response Elements* (*PRE*s) are required to repress some developmental target genes, and are widespread in genomes, suggesting that they broadly affect developmental programs. While *PRE*s in transgenes can nucleate trimethylation on lysine 27 of the histone H3 tail (H3K27me3), none have been demonstrated to be necessary at endogenous chromatin domains. This failure is thought to be due to the fact that most endogenous H3K27me3 domains contain many *PRE*s, and individual *PRE*s may be redundant. In contrast to these ideas, we show here that *PRE*s near the wing selector gene *vestigial* have distinctive roles at their endogenous locus, even though both *PRE*s are repressors in transgenes. First, a *PRE* near the promoter is required for *vestigial* activation and not for repression. Second, only the distal *PRE* contributes to H3K27me3, but even removal of both *PRE*s does not eliminate H3K27me3 across the *vestigial* domain. Thus, endogenous chromatin domains appear to be intrinsically marked by H3K27me3, and *PRE*s appear required to enhance this chromatin modification to high levels at inactive genes.

**Author summary:** Eukaryotic genes are packaged in chromatin, and their transcription relies on activators that recruit RNA polymerases and on repressive factors. Some of these factors factors that modulate chromatin structure to promote or inhibit gene transcription. In multicellular organisms, cell types have distinct patterns of gene expression, and these patterns are controlled by by the expression of cell-type-specific transcription factors and by modulating chromatin structure. The Polycomb system is one major system for the chromatin-mediated silencing of developmental gene expression, where a histone methyltransferase marks extended chromatin domains with trimethylation of lysine-27 of the histone H3 tail (H3K27me3) and forms repressed chromatin. In Drosophila, repressive regulatory elements called *Polycomb Response Elements* (*PRE*s) are thought to nucleate histone methyltransferase binding which then spreads across these domains. In this study, we demonstrate that two *PRE*s near the developmental *vestigial* gene have distinct and separable effects on gene activation and chromatin structure. Both *PRE*s are functional repressors in transgenes, but the *PRE* located near the *vestigial* promoter is required for gene transcription. This *PRE* has no effect on histone methylation of the domain. The second *PRE*, located in the middle of the chromatin domain is required for high-level H3K27me3 of the domain, and this methylation is not required to refine *vestigial* gene expression. Strikingly, a significant chromatin methylation remains when both *PRE*s are deleted. Our findings imply that *PRE*s near promoters may often play activating roles in gene expression in the Drosophila genome. We suggest that some domains of H3K27me3 occur in regions that lack histone acetylation, and has little consequence for correctly patterning gene expression.

## Introduction

The patterns of chromatin histone modifications differ between cell types, reflecting the activity of genes for developmental programs. Tri-methylation of the lysine-27 residue of histone H3 (H3K27me3) typically marks extended chromatin domains, leading to chromatin compaction and epigenetic gene silencing that is maintained as cells differentiate [1,2]. Histone methylation is thought to be initiated at discrete regulatory elements called *Polycomb Response Elements* (*PRE*s) within domains. These elements bind multiple DNA-binding factors, recruiting the PRC1 and PRC2 complexes, including the Polycomb chromatin factor and the E(z) histone methyltransferase, respectively [3,4]. Transgenes carrying *PRE*s are sufficient to silence reporter genes and to nucleate new H3K27me3 domains [5–7]. However, the function of *PRE*s in their endogenous domains is less clear. Deletion of *PRE*s from the homeobox gene cluster *BX-C* have limited defects in gene silencing [8–10], but no reduction of histone methylation of this domain. While multiple *PRE*s within the *BX-C* domain may be redundant, deletion of all mapped *PRE*s near the *invected* and *engrailed* genes have no effect on methylation of the locus, and it remains unknown how histone methylation is maintained [11].

Genomic mapping has identified regions where both the PRC1 and PRC2 complexes colocalize, and regions where each complex is found separately. Only about one-half of all Polycomb binding sites are within H3K27me3 domains, and thousands of additional sites are located near the promoters of active genes [12,13], where they may modulate gene expression by reducing release of RNAPII from paused promoters. Here, we characterize the *in vivo* roles of two *PRE*s near the *vestigial* gene. While these two *PRE*s are silencers in transgene assays, targeted mutations reveal that the promoter *PRE* is required for full gene expression. Using an new efficient method for genomic mapping of chromatin factors, we demonstrate that methylation across the domain remains in the absence of both *PRE*s. Our results reveal that *PRE*s stimulate but are not necessary for domain methylation.

## Results

### Profiling domains and regulatory elements in dissected tissues

To profile chromatin domains in different tissues, we adapted a chromatin mapping strategy that tethers micrococcal nuclease at factor binding sites. In the CUT&RUN procedure [14], unfixed cells are soaked with a factor-specific antibody, which binds to chromatin. Next, a protein-A-micrococcal nuclease (pA-MNase) fusion protein is soaked in, binding to the chromatin-bound antibody. Activation of the tethered MNase by adding calcium then cleaves exposed DNA around the binding sites of the targeted factor. As MNase cleaves at exposed DNA, the tethered nuclease typically releases both the bound factor and adjacent nucleosomes, and these particles can be distinguished by the length of the cleaved DNA (**Figure 1A**) [14]. In principle, tethered pA-MNase to a domain marked with a histone modification should similarly cleave apart all the nucleosomes of the domain, but also release all factor-bound DNA within the domain. Profiling the small fragment size class would then reveal regulatory elements within the domain as clusters of small reads (**Figure 1B**).

**Figure 1.**
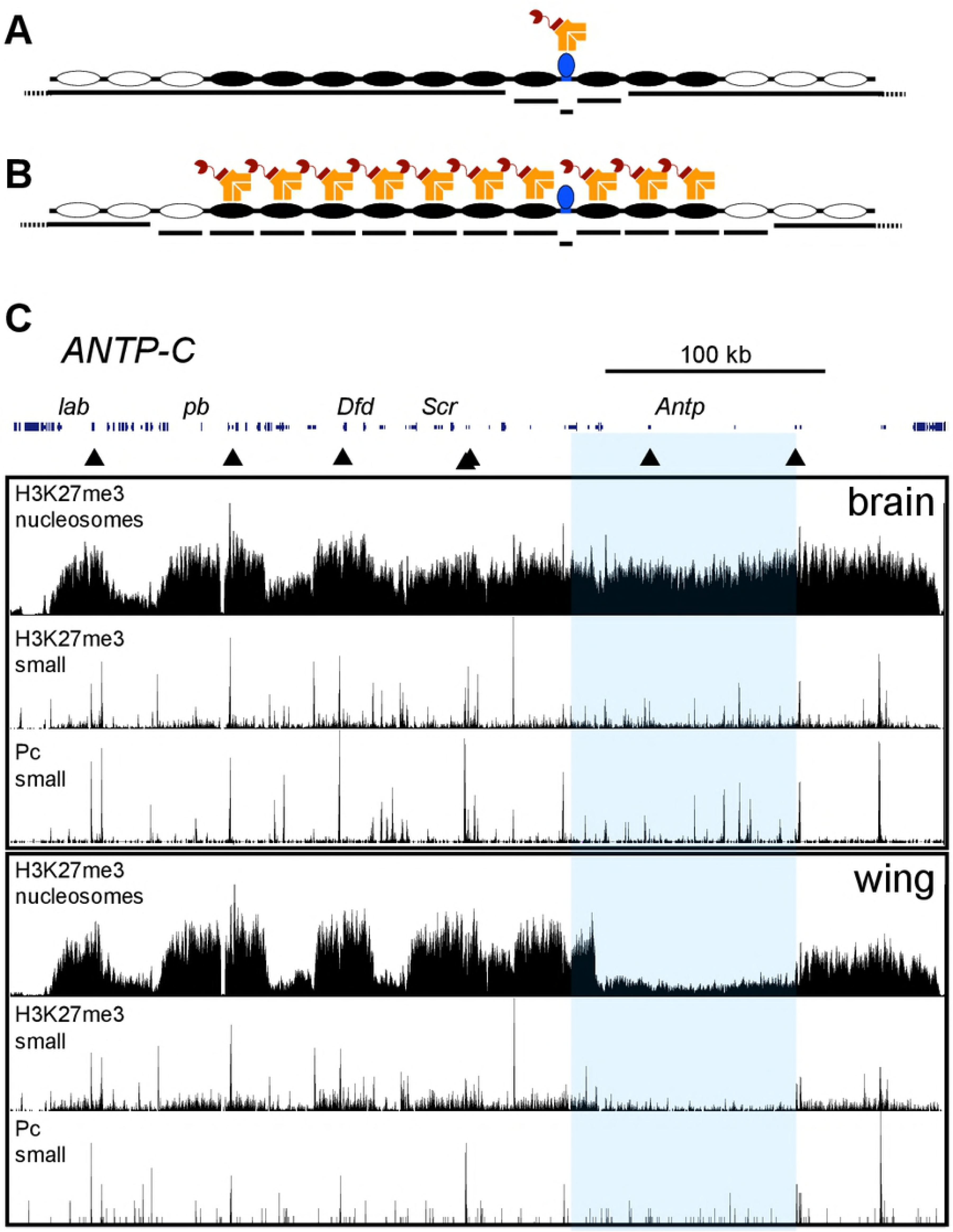
CUT&RUN profiling detects chromatin domains and embedded factor binding sites. (A) In CUT&RUN profiling for transcription factors, an antibody (yellow) is used to tether a pA-MNase fusion protein (red) to a bound transcription factor (blue). Activation of the nuclease cleaves exposed chromatin near the tethering site, producing small (20-120 bp) DNA fragments from the bound site and longer (>120 bp) fragments from adjacent nucleosomes. High-molecular weight DNA is left behind. Locations and lengths of fragments are determined by paired-end deep sequencing. (B) In CUT&RUN profiling of histone modifications, antibody tethering coats the chromatin domain with pA-MNase. Activation of the nuclease cleaves between nucleosomes and bound transcription factors within the domain, producing nucleosome-length and small DNA fragments, respectively. (C) Profiles of the 425 kb *ANTP-Complex* (chr3R:6,641,613-7,066,612 dm6) in larval brain and wing imaginal disc samples. The segment identity genes are labeled, and their promoters are marked by arrowheads. Landscapes for H3K27me3-tethered CUT&RUN are shown for nucleosome-length (120-710 bp) and small (20-120 bp) fragments, and for Polycomb-tethered CUT&RUN small fragments. The region including the *Antp* gene is highlighted in blue; this gene is silent in brain samples and transcribed in wing imaginal discs. *Antp* is packaged with methylated histones in brain samples, but methylation decreases across most of the *Antp* gene in wing discs. Multiple Polycomb-bound sites in this interval are present in brain samples, but disappear in wing samples.

CUT&RUN obviates the need to work with chromatin preparations or to optimize affinity recovery of chromatin particles, and works efficiently with small numbers of cells [15,16]. To implement CUT&RUN for tissue samples, we simply dissected brains and wing imaginal discs from 10 larvae, lightly permeabilized the whole tissues with digitonin, and sequentially incubated the tissues with antibody to H3K27me3 and then with pA-MNase. MNase was then activated and finally the cleaved DNA was isolated, subjected to Illumina paired-end sequencing, and mapped to the Drosophila dm6 genome assembly. We similarly mapped the Polycomb protein, which binds at *Polycomb Response Elements* (*PRE*s). We separated reads by length to distinguish nucleosomes (reads 120-710 bp) from factor-bound sites (small reads 20-120 bp in length).

H3K27me3 domains in larval tissues have been previously mapped by Chromatin Immunoprecipitation [17]. Profiles of H3K27me3 distribution generated by CUT&RUN using substantially less material were similar (**Figure 1C**). Both methods reveal changes in chromatin methylation that correspond to tissue-specific changes in gene expression. For example, the *ANTP-COMPLEX* cluster of homeobox genes are encompassed in a H3K27me3 domain in larval brains, consistent with the predominant silencing of this cluster in this tissue [18] (**Figure 1C**). In contrast, in wing imaginal discs where *Antp* is transcribed, chromatin over most of this gene is depleted for H3K27me3 (**FIgure 1C**). Interestingly, some histone methylation remains across the 3’ exons of *Antp*, implying that RNA polymerase II elongates through methylated chromatin during transcription.

CUT&RUN for the H3K27me3 modification also releases small DNA fragments, and these map in clusters within H3K27me3 domains (**Figure 1C**). In brain samples, eleven major clusters are apparent in the *ANTP-C* domain, and seven of these correspond to major Polycomb-bound sites, implying that these are *PRE*s. The other clusters correspond to known insulators and to gene promoters in the domain, implying that factors are bound there even though the genes are not transcribed. The remaining clusters presumably correspond to other regulatory elements. These landscapes highlight the utility of using small fragments within a histone modification domain to map regulatory elements.

There are striking changes in the chromatin landscape between tissues. In brain samples where the *Antp* gene is silenced and methylated, there are seven moderate H3K27me3-released small fragment clusters within the gene, and these show moderate Polycomb-binding (**Figure 1C**). In wing disc samples where the *Antp* gene is active and histone methylation is reduced, six of the small-fragment clusters disappear with an accompanying reduction in Polycomb signal in this region. Thus, activation of the *Antp* gene is accompanied by both the loss of the H3K27me3 modification and the loss of Polycomb binding at *PRE*s.

### The *vestigial* domain contains two *Polycomb Response Elements*

The multiple genes and regulatory elements within large H3K27me3 domains like *ANTP-C* make genetic analysis difficult. To define the effects of *PRE*s on endogenous target genes, we characterized the chromatin structure and expression of the developmental selector gene *vestigial* (*vg*), which is the only gene in a 32 kb H3K27me3 chromatin domain (**Figure 2A, top**). The *vg* gene is inactive in brain tissues, and is expressed only in the pouch of wing imaginal discs. Profiling of histone methylation reveals a somewhat reduced signal across the domain in wing disc samples (**Figure 2A, middle**), but these samples are a heterogeneous for *vg* expression. We therefore used the ‘Quadrant’ enhancer from the *vg* gene [19] to drive expression of GFP in the wing pouch, and used FACS to isolate GFP-positive cells. In these cells, histone methylation across the *vg* locus is greatly reduced (**Figure 2A, bottom**). Thus, histone methylation in this domain corresponds to the expression state of the *vg* gene.

**Figure 2.**
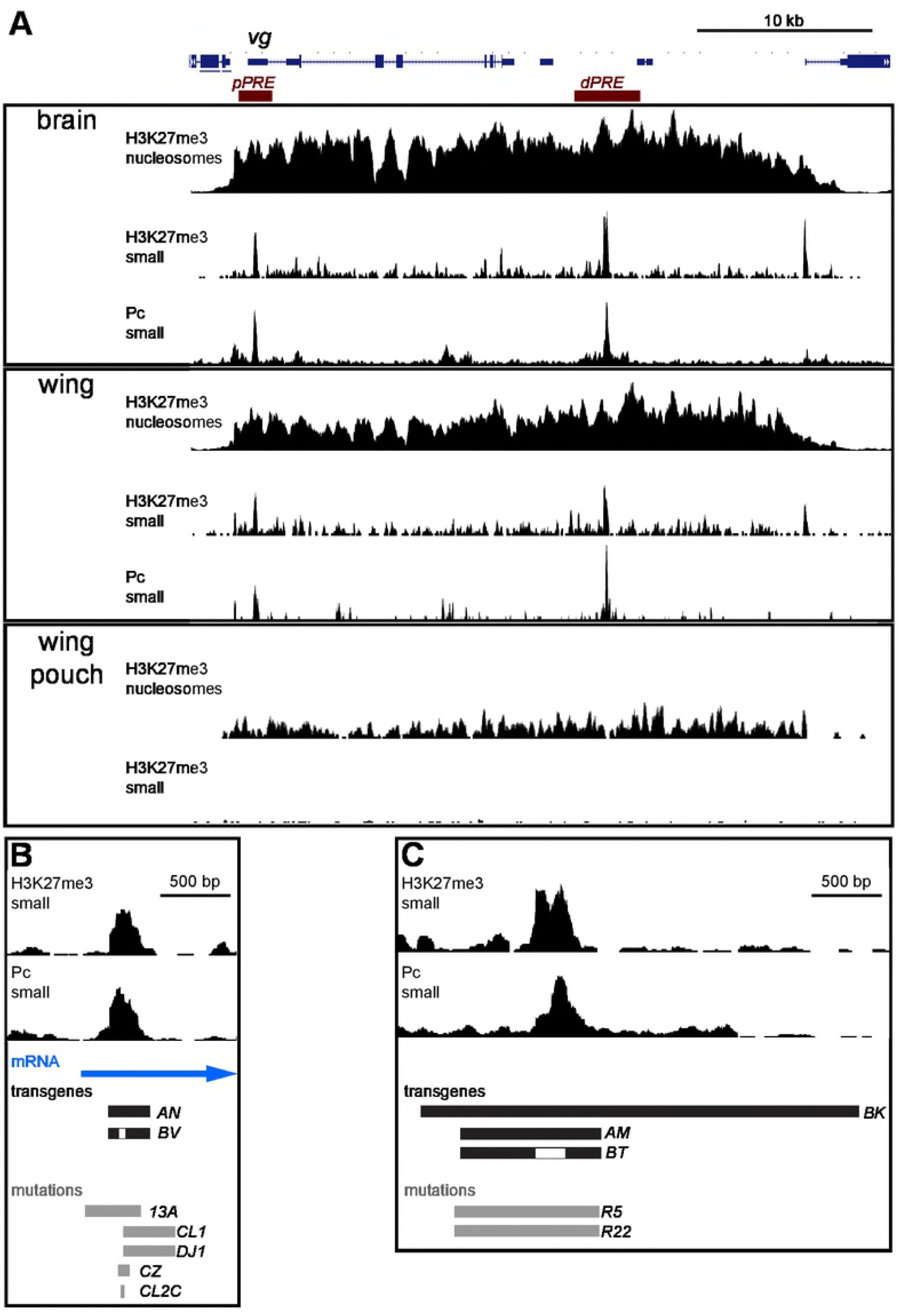
Chromatin features of the *vestigial* locus. (A) The *vestigial* wing-determination gene is included in a 32 kb domain of histone H3K27-trimethylation in larval brain and wing imaginal disc samples, but methylation is greatly reduced in FACS-isolated *vg*-expressing wing disc pouch cells. Tracks for nucleosome-sized (120-710 bp) and small (20-120 bp) fragments after H3K27me3-tethered CUT&RUN are shown, and for small fragments after Polycomb-tethered CUT&RUN. The major two putative *PRE*s (red boxes) have been previously mapped. (B-C) Molecular features of the two *PRE*s, showing small fragments after H3K27me3-tethered CUT&RUN, and after Polycomb-tethered CUT&RUN. Regions included in transgene constructs (black), and removed in mutations of the endogenous locus (grey) are marked. (B) A 2 kb segment around the *proximal PRE* (*pPRE*) (chr2R:12,883,995-12,885,995 dm6) with a Polycomb-bound site located +300 – +400 bp from the *vg* TSS within the *vg* transcribed region (blue). Constructed deletions remove the Polycomb-bound site (the *vg*^*CL1*^, *vg*^*13A*^, and *vg*^*DJ1*^ alleles), or the peak of the Polycomb-bound site (the *vg*^*CZ*^ allele). The *vg*^*CL2C*^ allele is a C->T substitution within the major Polycomb-bound site (chr2R:12,884497 dm6). (C) A 4 kb segment around the *distal PRE* (*dPRE*) (chr2R:12902222-12906221 dm6) with one Polycomb-bound site. Two engineered deletions (the *vg*^*R5*^ and *vg*^*R22*^ alleles) remove the site.

Two potential *PRE*s within this domain have been localized by sequence motifs [20] and by chromatin profiling [12,13,21]. These two *PRE*s coincide with small fragment clusters from H3K27me3-tethered CUT&RUN (**Figure 2A**). Further, mapping by CUT&RUN shows that Polycomb is bound at both of these sites (**Figure 2A**). The first site, which we term the *proximal PRE* (*pPRE*), is located 300 bp downstream of the mapped Transcriptional Start Site (TSS) of the *vg* gene [22]. The second *distal PRE* (*dPRE*) is located ~25 kb downstream (**Figure 2B,C**). Although other clusters of small fragments on each edge of the H3K27me3 domain might be minor *PRE*s, we first focused on the two major ones using transgenes and using mutagenesis of the endogenous locus.

Previous studies showed that sequences including the *dPRE* silence transgene reporter genes [23]. To test if the *pPRE* is also a silencing element, we tested genomic fragments for pairing-sensitive silencing (PSS), a diagnostic feature of *PRE*s where they silence adjacent reporter genes in homozygous animals [24]. Indeed, a transgene containing 300 bp including the *pPRE* and a *mini-w+* reporter gene integrated at the PhiC31 landing site shows strong PSS, as does a transgene containing the 3.1 kb *dPRE* identified in previous reports [23] (**Figure 3**). Thus, both of the putative *PRE*s in the *vg* domain appear to be functional silencers. These *PRE*s can interact with each other, as animals heterozygous for a *pPRE* transgene in one landing site and a *dPRE* transgene on the homolog also show PSS (**Figure 3**).

H3K27me3 CUT&RUN defined a 200 bp segment where Polycomb binds near the *vg* promoter. We used high-resolution mapping by native ChIP in Drosophila S2 cells to precisely define binding sites for three juxtaposed Polycomb-bound sites, one of which is also bound by the Pleiohomeotic (PHO) transcription factor (**Supplementary Figure 1**). Deletion of this site from the *pPRE* transgene alleviates PSS (**Figure 3**), thus this sequence is required for reporter silencing. We analyzed the *dPRE* similarly. We found that while a transgene including the *dPRE* induces PSS, deletion of the Polycomb-bound site within the *dPRE* alleviates this silencing (**Figure 3**). We conclude that the Polycomb-bound sites in both the *pPRE* and *dPRE* elements are required for transgene silencing.

**Figure 3.**
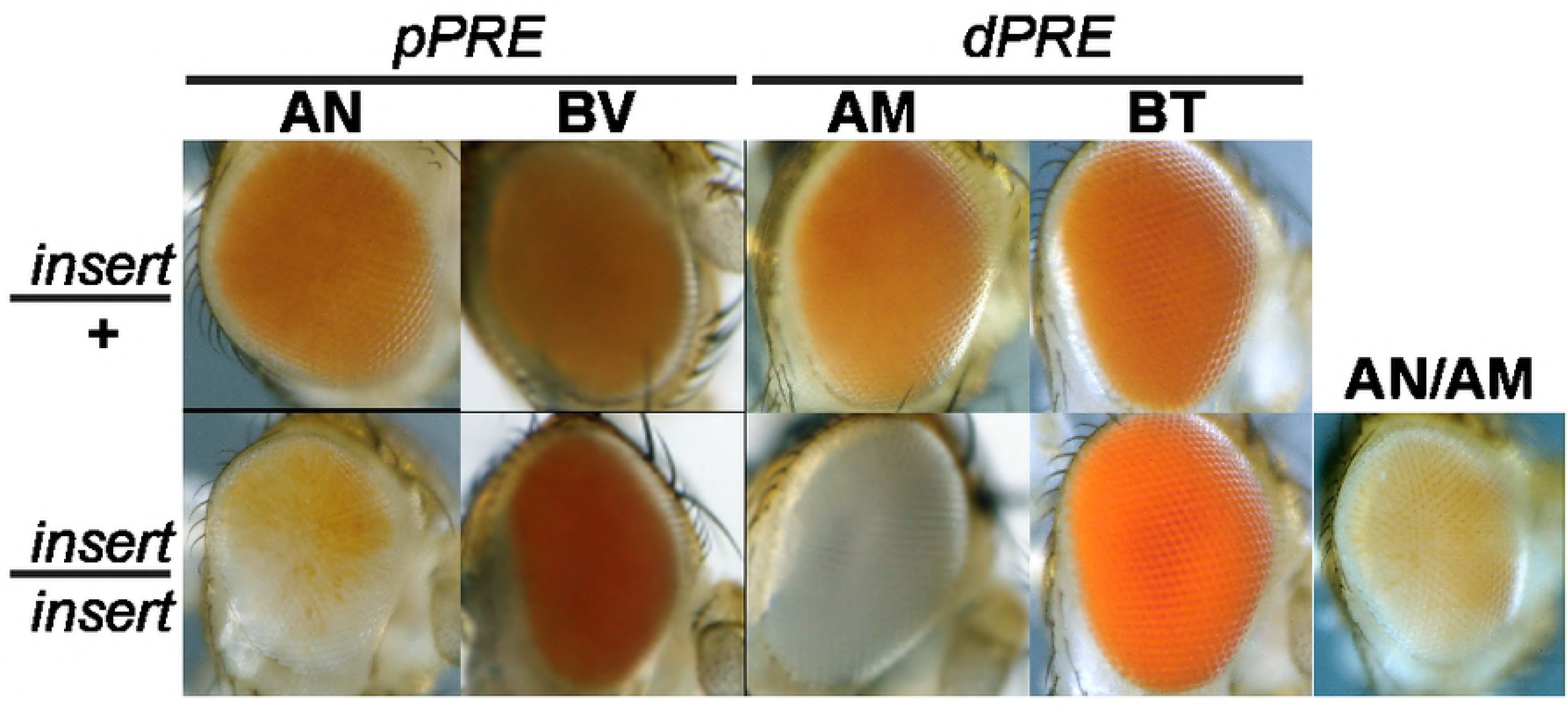
The *vestigial PRE*s mediate silencing of reporter genes. Position-Sensitive silencing (PSS) in the adult eyes of animals carrying *PRE* fragments next to a *mini-w+* reporter. Fragments from the *vg pPRE* and *dPRE* are indicated in Figure 2, and inserted in the same phiC31 landing site. Reduced *mini-w+* expression in animals homozygous for an insertion indicates silencing mediated by a *PRE*-containing fragment.

### Silencing of the *vg* domain requires H3K27 methylation

To measure silencing at the endogenous *vg* locus, we integrated reporter genes by gene-targeting near the *pPRE* and near the *dPRE*. The promoter of the *engrailed* gene is active in the posterior half of the wing imaginal disc [25], including part of the expression domain of *vg* in the wing pouch. We used an *engrailed-GAL4* (*en-GAL4*) transgene [26] to drive expression of GAL4-dependent *UAS-YFP* and *UAS-RFP* reporters in the wing disc. Control reporter gene insertions produce RFP and YFP throughout the posterior half of the wing disc (**Figure 4A**). In contrast, *UAS-YFP* reporters inserted in the *vg* domain are silenced throughout most of the wing disc, with reduced expression only within the posterior part of the wing pouch (**Figure 4A**). Insertions near the *dPRE* show similar reduced expression in the wing pouch and silencing in the rest of the wing disc. This implies that the *vg* domain is packaged in repressed chromatin in most of the wing disc, but derepressed in the wing pouch where the *vg* gene is transcribed. This is consistent with the change in histone methylation in wing disc pouch cells (**Figure 2A**).

**Figure 4.**
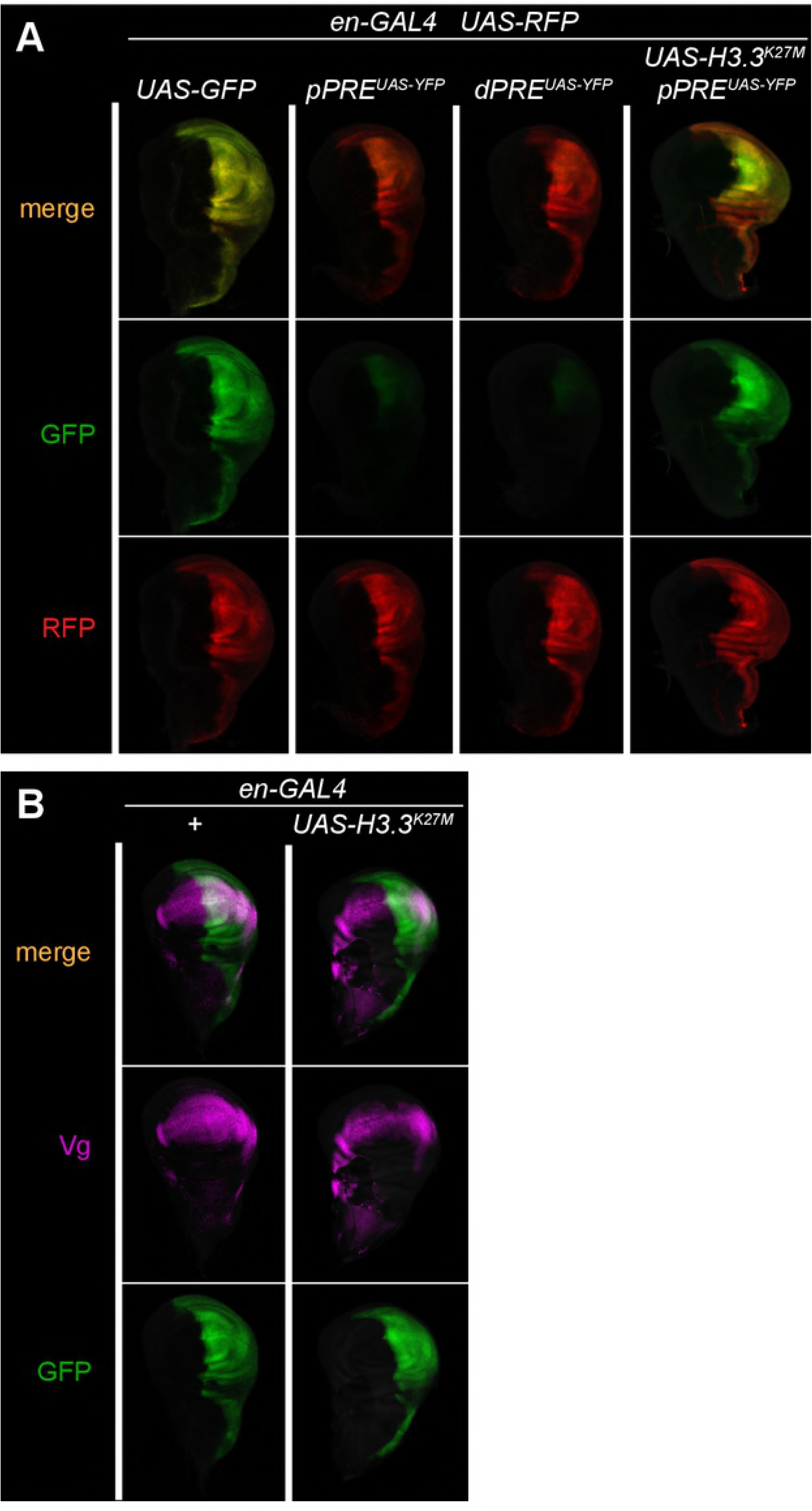
The *vestigial PRE*s mediate silencing of reporter genes. (A) Fluorescent protein expression in wing imaginal discs. Each animal carries the *en-GAL4* driver, and a *UAS-RFP* control reporter (red). Expression of *UAS-GFP* and *UAS-YFP* is in green. Control insertions outside of Polycomb-regulated domains shows high expression of GFP where *en-GAL4* is expressed. *UAS-YFP* insertions near the *vg pPRE* or *dPRE* show strong silencing, except in the wing pouch where *vg* is normally expressed. Silencing is eliminated by expression of a mutant H3.3^K27M^ histone in the posterior half of the wing disc. (B) The *vg* gene is not derepressed by the mutant H3.3^K27M^ histone. Expression of GAL4 was visualized with *UAS-GFP* (green), and the Vg protein by antibody staining (magenta).

We confirmed that silencing in the wing disc is mediated by chromatin by expressing a dominant-negative H3.3^K27M^ mutant histone to reduce chromatin levels of H3K27me3 [27] in the posterior half of the wing disc. Indeed, expression of the mutant histone derepressed the GFP reporter gene throughout the posterior half of the wing disc (**Figure 4A**). This demonstrates that the *vg* domain is in two chromatin states in the wing imaginal disc: a silenced configuration, and a derepressed configuration in wing pouch cells where *vg* is normally expressed. However, expression of the mutant histone does not derepress expression of the *vg* gene itself (**Figure 4B**). Thus, the silenced configuration appears to only affect the inserted reporter gene.

### The *proximal PRE* activates the *vestigial* gene

To define the function of each *PRE* for the *vg* gene, we deleted each element from the endogenous locus (see Methods). Precise breakpoints for each of the recovered deletions were determined by Sanger sequencing, and tested against each other and against previously characterized *vg* alleles (**Figure 2B,C**; **Supplementary Table 1**; **Supplementary Table 2**). Ectopic expression of *vg* converts legs and eyes into wing-like structures [28]; thus deletion of silencing elements should derepress the *vg* gene and transform non-wing tissues. However, we observed no such transformations in animals homozygous for deletion of the *pPRE* or of the *dPRE*. Deletion of both *PRE*s from the *vg* domain did not transform non-wing tissues, implying there is no role for Polycomb silencing in limiting normal *vg* expression. Surprisingly, deletions of the *pPRE* reduce expression of the *vg* gene. Animals carrying the *vg*^*CL1*^ or *vg*^*13A*^ deletions in combination with a null allele showed severe loss of wing tissue (**Figure 5A**). The adult wing blade differentiates from the pouch of the wing imaginal disc where *vg* is expressed (**Figure 5B**); in wing imaginal discs from animals with a *pPRE* deletion no Vestigial protein is detectable, the wing pouch is reduced, and the central stripe of Wingless expression marking the future wing blade margin is absent (**Figure 5B**). Thus, *pPRE* deletions are severe loss-of-function alleles. A smaller deletion within the Polycomb-bound segment of the *pPRE* (the *vg*^*CZ*^ allele) shows a partial loss of the distal wing blade; thus, this allele has reduced *vg* expression (**Figure 5A**). In contrast, animals carrying the *vg*^*R5*^ deletion of the *dPRE* show no defect in wing morphology (**Figure 5A**), demonstrating that this *PRE* has no positive or negative role in normal *vestigial* expression. Animals lacking both *PRE*s are wingless like the *pPRE* single mutant (**Figure 5A**).

**Figure 5.**
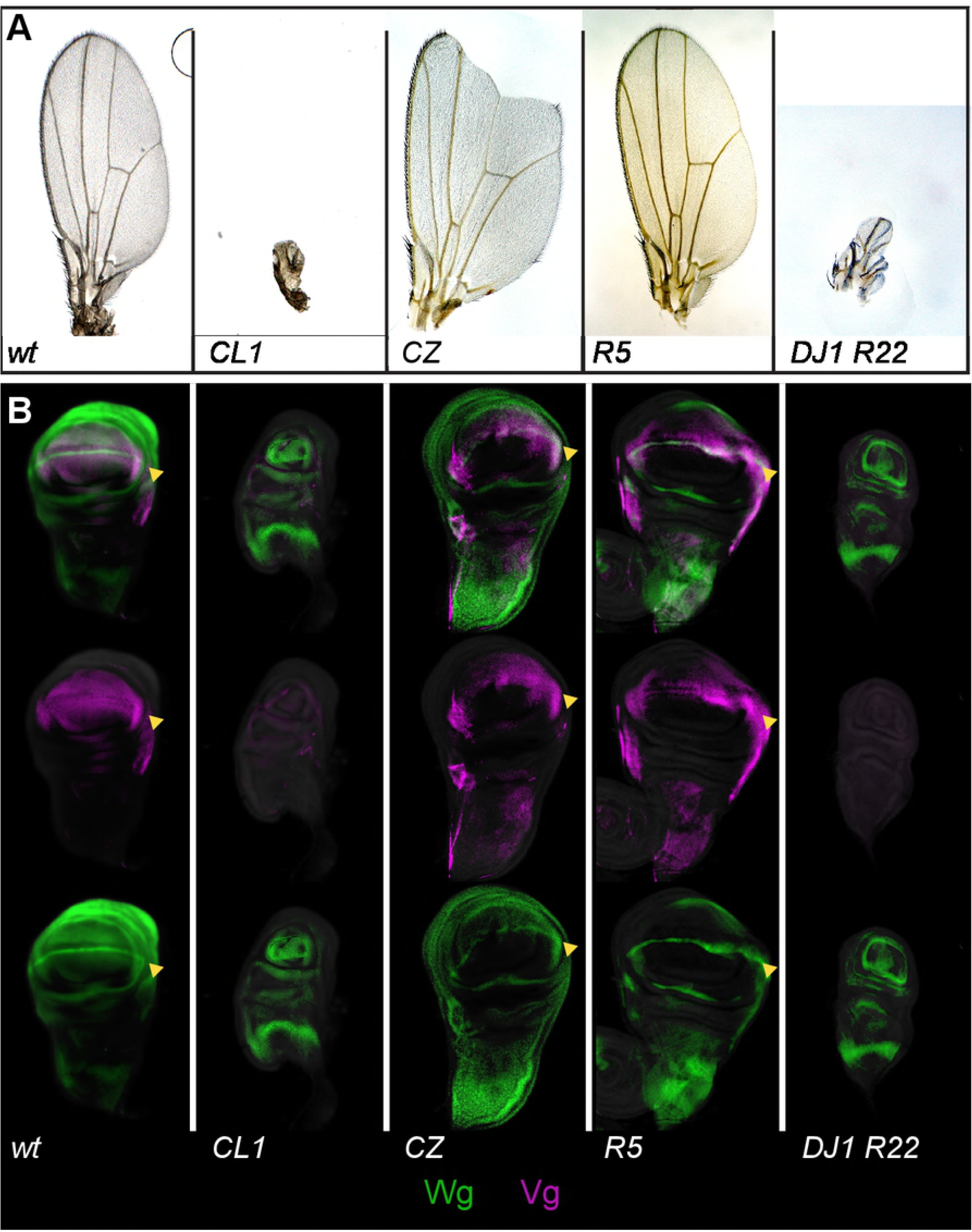
The *pPRE* and Polycomb are required for *vestigial* activation. (A) Wings from adults with the indicated genotypes. New alleles were heterozygous with the *vg*^*nw*^ null allele to show expression of each allele. (B) Wing imaginal discs stained with anti-Vg (magenta) and anti-Wg (green) antisera. In wildtype, a ring of Wg circles the wing pouch, and a stripe of Wg across the wing pouch marks the future adult wing blade margin (margin indicated by a yellow arrowhead). Vg protein is produced throughout the wing pouch. In *vg*^*CL1*^/*vg*^*nw*^ animals no Vg protein is detected, the wing pouch is greatly reduced, and the future wing margin is missing. In *vg*^*CZ*^/*vg*^*nw*^ animals expression of Vg is reduced, and gaps are apparent in the wing margin Wg stripe, anticipating the wing margin notches in adults of this genotype. Both Vg and Wg expression patterns appear normal in the *dPRE* deletion *vg*^*R5*^. Finally, the double *PRE* deletion *vg*^*DJ1 R22*^ resembles the single *pPRE* deletion, with loss of Vg expression and of the margin stripe of Wg.

The opposite effects of the *pPRE* on the *vg* gene compared to its effect on reporter transgenes suggests that activation of the *vg* promoter may requires Polycomb factors. Indeed, we recovered one point mutation in the *pPRE* that supports this idea. The *vg*^*CL2C*^ allele is a C-to-T substitution in the *pPRE* that lies precisely at the center of the major Polycomb-bound site, in a sequence similar to consensus motifs for Sp1 transcription factors, which have been implicated in Polycomb targeting [29] (**Figure 2B**; **Supplementary Figure 1**). This allele causes moderate notching of the adult wing, implying that binding at this site is required for full *vg* activation (**Figure 6A**). We tested if Polycomb binding at this site represses or promotes *vg* expression by combining the *vg*^*CL2C*^ allele with null mutations in *Polycomb*, which partially derepresses silenced genes [30]. While animals heterozygous for the *vg*^*CL2C*^ allele have no phenotype, in combination with a heterozygous *Pc*^*3*^ allele animals have crumpled wings with small notches in the distal wing margin (**Figure 6A**). Wing defects are more severe in *vg*^*CL2C*^/*vg*^*nw*^; *Pc*^*3*^/+ animals. Mutations in the RING1b homolog *Sce* similarly enhance the phenotype of *vg*^*CL2C*^/*vg*^*nw*^ animals (**Figure 6A**). These effects imply that a PRC1 complex including Polycomb and Sce bound at the *pPRE* has a positive effect on *vg* expression.

**Figure 6.**
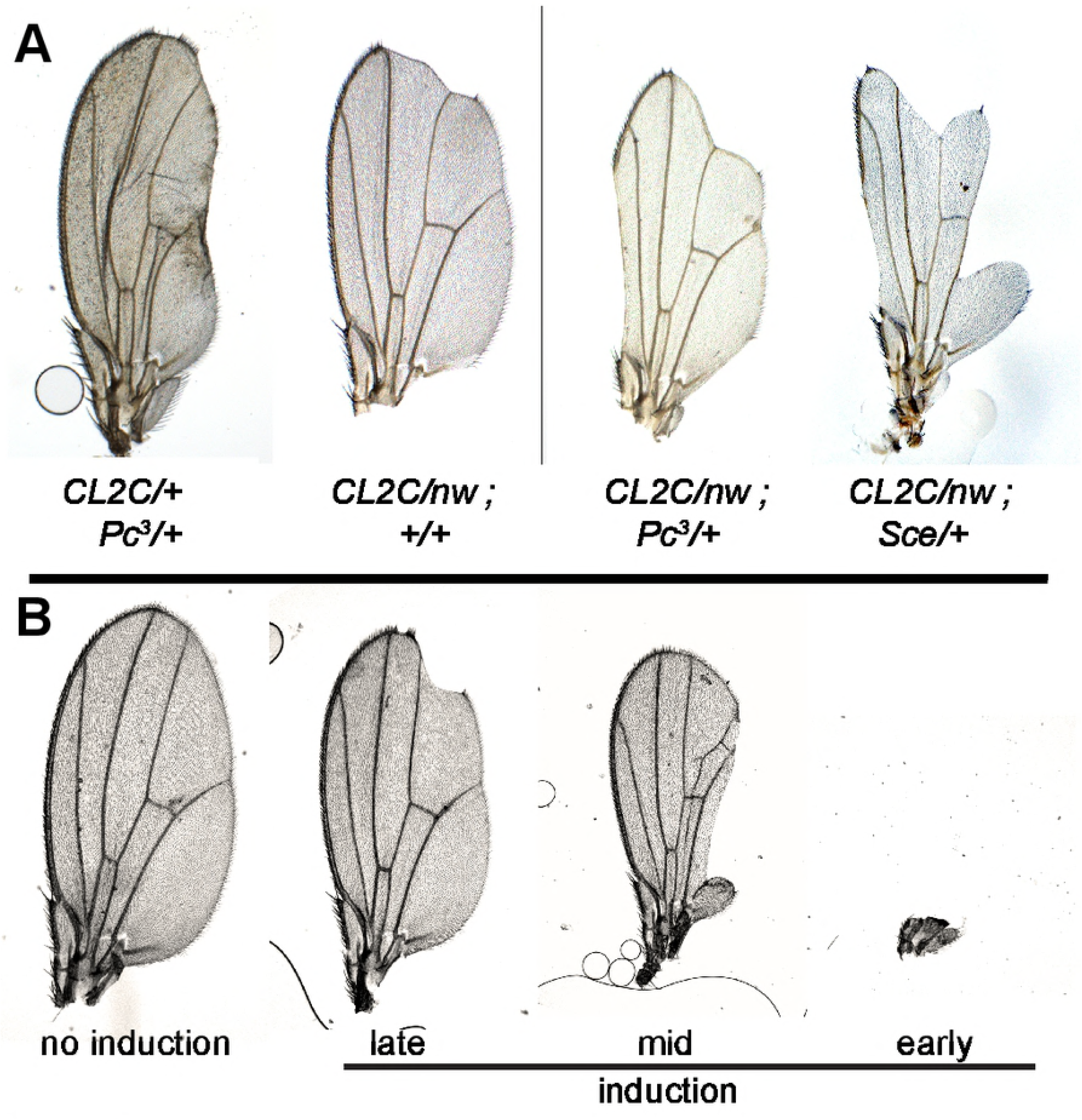
The *pPRE* is required in active cells. (A) Wings from adults with the indicated genotypes. The *pPRE* substitution mutation *vg*^*CL2C*^ is a recessive mutation, but reduction of Polycomb results in bowed and crumpled wings. *Polycomb* or *Sce* mutations enhance the wing defects of *vg*^*CL2C*^/*vg*^*nw*^ animals. (B) Wing morphologies after *FLP* induction during embryonic and larval development. Most wings show no defects in morphology. Occasional adults that experienced FLP induction during development show notches in the wing margin (induced in late larvae), deletion of portions of the wing (induced in early larvae), or complete loss of the wing (induced in embryos).

It is unusual for PRC1 to be implicated in transcriptional activation. In one case in the mouse midbrain, Polycomb is required to bring enhancers to the *meis2* gene promoter before *meis2* is expressed, but then is not required after induction [31]. To test if the *vg pPRE* is similarly required before activation of the *vg* gene or if the *pPRE* is required in cells expressing *vg*, we generated cells homozygous for a *pPRE* deletion from heterozygous cells by *FLP* recombinase-mediated mitotic recombination at different times in development [32]. The *pPRE/+* heterozygous animals have no wing defects, but *FLP* expression produces animals with a range of defects in the wing blade, ranging from notches in the wing margin to complete loss of one wing (**Figure 6B**), implying that *pPRE* mutant clones lose *vg* expression whenever they are induced. Together, these results indicate that a PRC1 complex bound at the *pPRE* is required to maintain expression of the *vg* gene.

### The *distal PRE* is required for chromatin methylation

In transgenes, a *PRE* is required to nucleate and maintain a Polycomb-regulated domain by recruiting PRC1 and PRC2 complexes [6,7]. We tested if histone methylation of the *vg* domain depends on the *pPRE* or on the *dPRE*. We normalized landscapes by the height of signals across the *ANTP-C* domain, and then compared the genomic region including the *vg* domain in wing imaginal discs from wildtype and *PRE* deletion mutants (**Figure 7, top**). H3K27me3 levels are moderately high across the *vg* domain in wildtype wing discs, which contain cells with and without *vestigial* expression. In animals lacking the *pPRE* only non-*vestigial*-expressing cells are present, and H3K27me3 levels across the domain are increased. In contrast, domain methylation is reduced to ~25% of wildtype levels in animals lacking the *dPRE*. Wing discs from animals with deletions of both *PRE*s show reduced – but not absent – levels of chromatin methylation across the *vg* domain, even though these discs contain only *vestigial* non-expressing cells. These results imply that only the *dPRE* is required for high-level methylation of the domain, but that a low level of methylation across the domain is independent of both *PRE*s.

**Figure 7.**
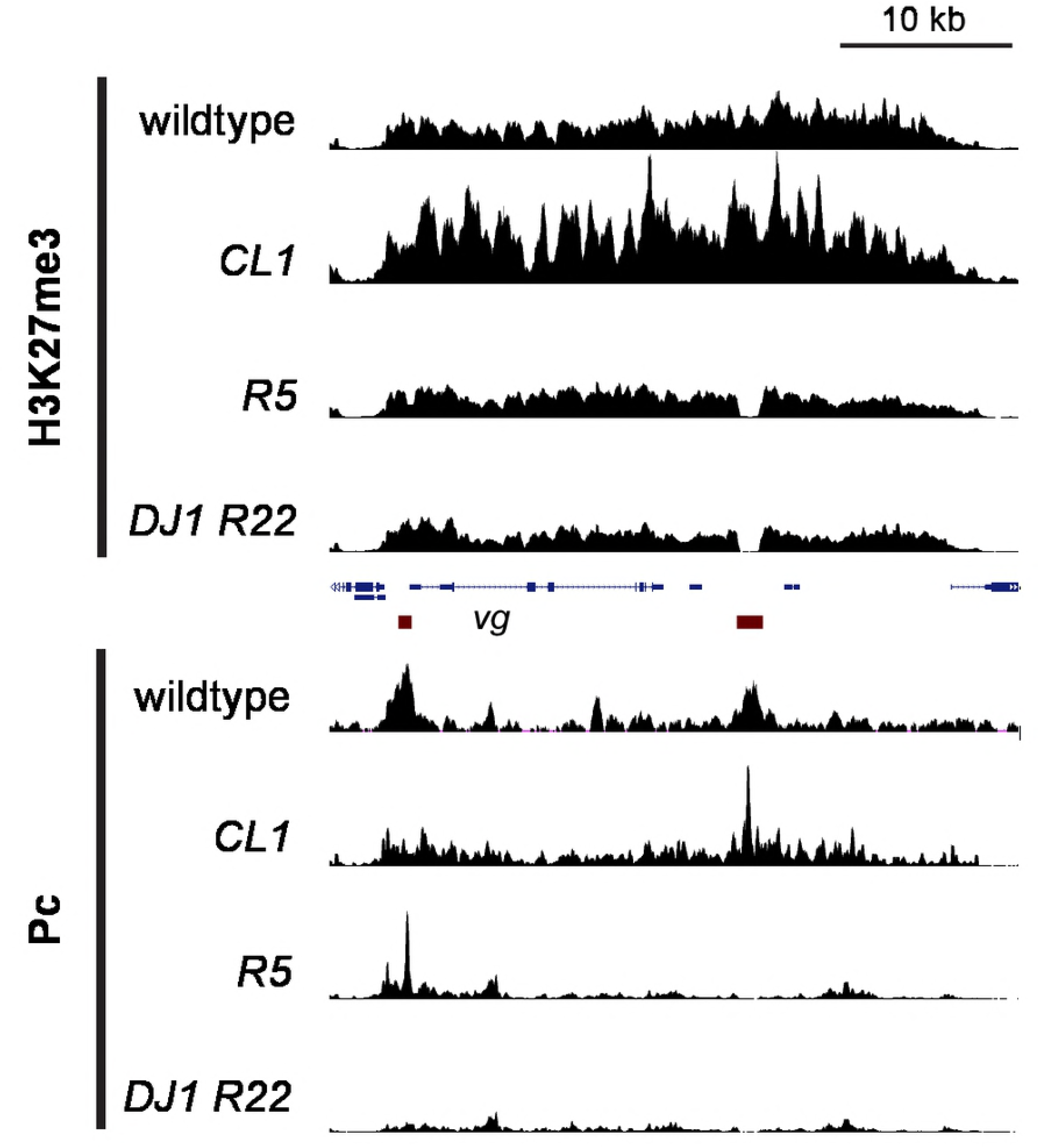
Histone methylation of the *vestigial* domain is reduced in *PRE* mutants. Wing imaginal discs from wildtype animals and from *dPRE* (*vg*^*R5*^) mutants are a mixture of *vg*-expressing and non-expressing cells, while discs from the *pPRE* deletion mutant (*vg*^*CL1*^) and the double mutant (*vg*^*DJ1 R22*^) lack the wing pouch and are thus only *vg*-non-expressing cells. Genes (blue) in the domain and the two *PRE*s (red) are indicated. H3K27me3 profiles (top 4 tracks) show moderate levels of domain methylation in wildtype discs, and high levels of H3K27me3 across the inactive *vg* domain in *vg*^*CL1*^ mutants. Methylation is reduced when the *dPRE* is deleted (*vg*^*R5*^), indicating that this *PRE* nucleates methylation of the domain. Discs from both *vg*^*R5*^ and the double *PRE* mutant *vg*^*DJ1 R22*^ have reduced but substantial methylation remaining across the *vg* domain. The Polycomb landscapes (bottom 4 tracks) across the *vg* domain shows two major peaks in wildtype wing imaginal discs, coinciding with the *pPRE* and the *dPRE*. No additional Polycomb signals are apparent in the *vg* domain in the *vg*^*DJ1 R22*^ *PRE* double mutant. Polycomb binding in all four genotypes was normalized to peak heights in the *ANTP-C* domain.

It is possible that minor or cryptic *PRE*s in a domain may direct histone methylation in *PRE* deletion mutants [11]. We therefore profiled Polycomb binding in wing discs from wildtype and *PRE* deletion mutants, normalizing landscapes to peak heights in the *ANTP-C* domain (**Figure 7, bottom**). The two major peaks of Polycomb binding in the *vg* domain coincide with the *pPRE* and the *dPRE*, and deletion of either of these eliminates only its peak. Notably, Polycomb is bound at the *pPRE* in *dPRE* deletion mutants, but this is not sufficient for high-level histone methylation of the domain. Further, when both *PRE*s are deleted there is no peak of Polycomb binding in the *vg* domain, indicating that there are no alternative or cryptic *PRE*s in the *vg* domain. Thus, a low level of histone methylation across silenced domains can be independent of any *PRE*.

## Discussion

The concept that Polycomb-repressed chromatin domains are nucleated at short factor-binding regulatory elements (*PRE*s) derives from the phenotypes of deletions within the homeobox clusters in Drosophila, where *PRE*s are required for silencing of these genes [33]. However, it has been difficult to determine if *PRE*s control histone modifications in these domains. While transgenes can confer H3K27me3 modification onto their insertion sites [6,34,35], the effects of deleting *PRE*s from endogenous domains has been more ambiguous. In part these ambiguities may be due to the difficulties of measuring changes in histone modifications with limited samples. Using the more efficient CUT&RUN method, we find that we can produce detailed chromatin profiles from small samples with high sensitivity, allowing characterization of specific tissues in mutant animals. CUT&RUN both details histone modification domains and reveals the locations of transcription factor binding sites within domains of histone modifications, providing a *de novo* method to map potential regulatory elements.

### *PRE*s are only partially responsible for domain methylation

The developmental gene *vestigial* is contained within chromatin that has all the features of a Polycomb chromatin domain, being marked by the H3K27me3 histone modification and silencing inserted reporter transgenes. While the two *PRE*s from this domain both act as silencers in transgenes, our results show that they have distinct roles at the endogenous locus: the proximal *PRE* affects *vg* expression, but has no effect on histone methylation of the domain, while the distal *PRE* increases histone methylation but has no effect on expression. A similar specialization of a *PRE* that causes histone methylation has been identified at the *dachshund* locus in Drosophila [36]. These effects support the idea that the E(z) H3K27 methyltransferase is recruited to nucleate histone methylation of the domain. In contrast, deletions of the known *PRE*s of the *engrailed* locus had no effect on domain methylation. It was suggested that additional cryptic or weak *PRE*s are present in the *engrailed* domain [11], but no such elements are apparent in our profiling of the *vg* domain. Non-coding RNAs have been proposed to target and modulate histone methylation in many organisms, and such RNAs have been identified in the *vg* domain [37]. However, these RNAs are deleted in our mutation of the distal *PRE*, and thus cannot be responsible for the residual methylation of the *vg* domain.

Active chromatin regions are often marked by histone acetylation, including acetylation of the H3 K27 residue, and this modification is antagonistic to H3K27 methylation [38]. In Drosophila cells, untargeted E(z) methyltransferase methylates ~50% of all nucleosomes to a dimethylated state [39]. Regions without acetylation may continue to accumulate methylation, and this background histone methylation may prime regions for more extensive methylation by *PRE*s. This implies that genomic patterns of histone methylation could then be determined in part by the pattern of histone acetylation, and some chromatin domains may not require *PRE*s for their specification, if they are defined by the absence of activating elements.

### A promoter *PRE* activates gene expression

*PRE* localization near promoters is a common feature of the Drosophila genome [12,40]. Such *PRE*s are well-positioned to regulate gene activity. The transition of initiating RNA polymerase II (RNAPII) to its elongating form is a major step for controlling the expression of developmental genes [41]. Inhibiting the elongation of RNAPII is one mechanism by which Polycomb can silence genes [42]. However, a number of developmental genes are now known to have little or no reliance on chromatin repression, despite having Polycomb-bound promoters [11,36,43]. This may result because Polycomb is recruited to chromatin by a number of ubiquitous transcription factors that are generally used at regulatory elements [1,4,13]. Such promiscuous binding results in many elements with chromatin features of Polycomb repression, but with limited effects on regulation of of many endogenous genes.

Polycomb has also been implicated in the regulation of active genes. These *PRE*s have been suggested to reduce transcriptional output [12], but loss of Polycomb can decrease elongating RNAPII at many active genes as well [40]. The *vg* gene is the first example of a promoter that requires a nearby *PRE* for activation during development. Perhaps a *PRE* can loop and interact with more distant silencing elements or with enhancers, and the effects of a *PRE* may then depend on what regulatory elements it brings together. Indeed, Some *PRE*s have been reported to loop together enhancers [31], and can switch between silencing and activating states [44]. Our observations that a Polycomb domain may simultaneously silence an inserted gene and activate the endogenous gene argues that promoters differ in their interactions with *PRE*s, and highlights a central aspect of Polycomb regulation, where silencing and activating functions must be integrated with developmentally-programed enhancers.

## Methods

### Fly strains

All crosses were performed at 25˚C. Transgenes, mutations and chromosomal rearrangements not detailed here are described in Flybase (http://www.flybase.org). The *vg*^*nw*^ allele is a deletion of the last two exons of the *vg* transcript, and so we used this as a standard null allele. New alleles of *vg* produced in this study are described in **Supplementary Table 1**.

#### Transgene integration

Genomic *PRE*-containing segments were PCR amplified from *Oregon-R* genomic DNA and cloned by Gibson assembly into the split-*mini-w+* vector pKC27mw [23] (a gift from L. Ringrose, Humboldt-Univerität zu Berlin). Deletions within these genomic segments were generated by PCR amplification of the plasmid. The genomic segments in all constructs was verified by sequencing. Plasmids were injected into *y* M[*vas-int.Dm*]ZH-2A *w*; *P*[*attP,y+,w*^*3*^]VIE-260B embryos by Bestgene Inc (Chino Hills, CA). This line contains two landing sites [45]; integrants at the 25C landing site were used in this study. A line with an integrated plasmid with no genomic fragment was used as a negative control. Finally, a line carrying the *vg* Quadrant enhancer (chr2R:12,986,173-12,896,970 dm6) upstream of GAL4 was used (named *vgQ-GAL4*.CO) was used to label cells in the developing pouch of the wing imaginal disc.

#### P *mutagenesis*

The *vg*^*21-3*^ allele carries a *P* element inserted at the *pPRE* (chr2R:12,884,623 dm6). The *P* element was mobilized by crossing to the *TMS, P*[Δ*2-3*]99B transposase source, mosaic males were crossed to *vg*^*nw*^/CyO females, and lines were established from single animals with reduced wing phenotypes. Each mutant was characterized by sequencing an amplicon spanning the *vg*^*21-3*^ insertion site.

#### CRISPR mutagenesis

Guide RNAs to direct Cas9 to the *pPRE* were cloned by overhang PCR into the pU6sgshort plasmid [46] (a gift from N. Perrimon, Harvard Medical School, Boston). Two plasmids with guide RNAs to either side of the *pPRE* were co-injected into *y M[nos-Cas9.P, w+]*ZH-2A *w* embryos by Bestgene Inc (Chino Hills, CA). These mosaic animals were mated to *w; vg*^*nw*^/*CyOG*, and *Cy*^+^ progeny with reduced wings were used to establish stocks.

#### Targeted integration and resolution

We used a two-step strategy (**Supplementary Figure 2**) to first create a tandem duplication of a deleted *PRE* and a marker gene next to the wildtype *PRE* at the endogenous locus, and then in a second step to delete the wildtype *PRE* and the marker gene, leaving the deleted *PRE* in the locus. To create the tandem duplication, we used CRISPR to induce a double-strand break at a genomic target site, which invades homology on an injected plasmid, integrating it into the chromosome. In the second step, we induced expression of the *ISce-I* site-specific endonuclease to cleave within the tandem duplication (similar to [47]), stimulating recombination between the repeated sequences and resolving to an unmarked locus with the mutant *PRE*.

To do so, we cloned a *UAS-YFP* transgene into pUC19, then cloned genomic segments including the *vestigial pPRE* (1.8 kb) or the *dPRE* (2.9 kb). Deletions within the genomic segments were generated by PCR amplification of the plasmid. Guide RNAs to direct Cas9 to the *pPRE* or to *dPRE* were cloned into pU6sgshort, and co-injected with a plasmid carrying homology on each side of a *PRE* into *y M[nos-Cas9.P, w+]*ZH-2A *w* or *y w*^*1118*^; *attP2{nos-Cas9}/TM6C,Sb Tb* embryos by Bestgene Inc (Chino Hills, CA). Injected mosaic animals were crossed to *w; P*[*GMR-GAL4*]D *P*[*UASRFP, w*^+^]3, and progeny with YFP-positive eyes were used to establish stocks and then the order of the duplicated segments was determined by PCR. Lines with a tandem duplication in the order (mutant *PRE*–UAS-YFP–*wildtype *PRE*) were crossed to *y w; P*[*HS-ISce-I, v*^+^]2B *Sco* / *S*^*2*^*CyO*, and progeny were heat-shocked to express the endonuclease. Mosaic *y w / Y; PRE*–UAS-YFP–PRE / P*[*HS-ISce-I, v*^+^]2B *Sco* males were crossed to *w; P*[*GMR-GAL4*]D *P*[*UASRFP, w*^+^]3, and Sco+ males with no YFP expression were used to establish stocks. Resolution of tandem duplications was confirmed by Sanger sequencing.

To construct a mutant chromosome deleted for both the *pPRE* and the *dPRE*, we injected two plasmids encoding guide RNAs targeting the *pPRE* into a *dPRE*–UAS-YFP–dPRE* tandem duplication stock, and recovered a *pPRE* deletion by screening for reduced wings. We then resolved the tandem duplication to delete the *dPRE*, producing a double mutant chromosome.

#### Generation of mutant clones

Vials with *P*[*hsFLP, ry*^+^]1 *y w; P*[*FRT-w*^*hs*^-*FRT*]G13 *vg*^*CL1*^ / *P*[*FRT-w*^*hs*^-*FRT*]G13 larvae were heat-shocked at 37˚ for 1 hour in a water bath to induce *FLP* expression and mitotic recombination. Adults were examined for defects in wing morphology.

#### Imaging wing imaginal discs

We dissected wing imaginal discs from late 3rd instar larvae and fixed them for 20 minutes in 4% formaldehyde/PBST (PBS with 0.1% triton-X100). For detection of proteins in wing discs, tissues were blocked with 10% goat serum/PBST, and incubated with anti-Vg (1:100 dilution, a gift from S Carroll, University of Wisconsin-Madison) and anti-Wg (1:200 dilution, clone 4D4, Developmental Studies Hybridoma Bank) antiserum at 4˚ overnight, and with fluorescently-labeled secondary antibodies (1:200 dilution, Jackson ImmunoResearch). All tissues were stained with 0.5 µg/mL DAPI/PBS, mounted in 80% glycerol on slides, and imaged by epifluorescence on an EVOS FL Auto 2 inverted microscope (Thermo Fisher Scientific) with a 10X objective.

#### Imaging adult wings

Wings were wetted in ethanol and mounted in 80% glycerol on slides, and photographed using a Sony digital camera mounted on a Nikon SMZ1500 stereomicroscope or with a 4X objective on an EVOS FL Auto 2 inverted microscope (Thermo Fisher Scientific).

#### Imaging adult eyes

Adult heads were mounted on steel pins, immersed in mineral oil, and imaged using a Sony digital camera mounted on a Nikon SMZ1500 stereomicroscope.

### Biological material for chromatin profiling

#### Drosophila cell culture

S2 cells were grown to mid-log-phase in HyClone Insect SFX media (GE). We used 1×10^6^ cells for each chromatin profiling experiment.

#### Intact larval brains and wing imaginal discs

We dissected tissues from 3rd instar larvae in Insect SFX media. We used ~10 larval brains and wing imaginal discs for each chromatin profiling experiment.

#### Isolated wing pouch cells

We dissociated and isolated cells from the vestigial-expressing portion of wing imaginal discs as described [48], with the following modifications. Approximately 200 wing imaginal discs from *vgQ-GAL4*.CO *P*[*UAS-GFP, w*^+^]T2 3rd instar larvae were dissected in SFX media and dissociated in Accutase (Sigma) for 1 hr at room temperature with occasional agitation. Accutase was blocked with 0.5 volumes of FBS. The solution was passed through a 40 µm filter and sorted for GFP fluorescence using 488 nm excitation and 530/30 nm emission filters on an BD Aria II Flow Cytometer using a flow rate of 3 (~20 µL/min) and a 100 µm nozzle. We used ~1×10^4^ cells for each chromatin profiling experiment.

### Chromatin profiling

#### CUT&RUN

We used an immuno-tethered strategy for profiling histone modifications and Polycomb binding in Drosophila cells. The CUT&RUN method uses an antibody to a specific chromatin epitope to tether a protein A fused to micrococcal nuclease (pAMN) at chromosomal binding sites within permeabilized cells [14]. The nuclease is activated by the addition of calcium, and cleaves DNA around binding sites. Cleaved DNA is isolated and subjected to paired-end Illumina sequencing to map the distribution of the chromatin epitope. We used primary antibodies to histone H3 trimethylation at lysine-27 (clone C36B11, Cell Signalling Technology, Inc), and to Drosophila Polycomb (a gift from G Cavalli, Université de Montpellier) We used a low-cell number version of CUT&RUN [15] for Drosophila S2 cells, intact larval tissues, and cells collected after FACS with the following modifications. S2 cells and cells collected after FACS were bound to BioMag Plus Concanavalin-A-conjugated magnetic beads (ConA beads, Polysciences, Inc), permeabilized in dbe+ buffer (20 mM HEPES pH 7.5, 150 mM NaCl, 0.9 mM spermidine, 2 mM EDTA, 0.1% BSA, 0.05% digitonin with Roche cOmplete protease inhibitor), cleaved in WashCa+ buffer buffer (20 mM HEPES pH 7.5, 150 mM NaCl, 0.9 mM spermidine, 0.1% BSA, 2 mM CaCl_2_ with Roche cOmplete protease inhibitor) at 0˚ for 30 minutes, and then DNA was recovered as described [15]. In some cases we collected S2 cells by centrifugation and cleaved chromatin for 5’ at room temperature. For intact larval brains and wing discs, we transferred tissues between buffers in in the wells of glass dissection dishes by pipetting with low-retention tips. Tissues were incubated with dbe+ buffer, incubated in primary antibodies overnight at 4˚ with gentle rocking, and incubated in pAMN/dbe+ for 1 hour at room temperature. After washing, tissues were transferred to eppendorfs with chilled WashCa+ buffer and incubated at 0˚ for 30’ for DNA cleavage, then stopped, treated with RNase A for 30 minutes at 37˚, and spun at 16,000 g for 5 minutes at 4˚ to separate supernatant and pellet fractions, and processed as described [15]. In all experiments we added 0.3 pg of yeast fragmented DNA as a spike-in normalization control for library preparation and sequencing [14]. Two replicates were performed for each experiment. In some experiments we coated dissected tissues with ConA beads and moved them between solutions using a magnet, and recovered cleaved DNA with Ampure XP beads (Beckman Coulter) immediately after protease treatment. A step-by-step protocol is posted (https://protocols.io/private/D6B0AD2DC1431A513994A2A05AC59CDA).

#### Library preparation, sequencing, and data processing

Libraries were prepared as described [14], with 14 cycles of PCR with 10 second extensions for enrichment of short DNA fragments. Libraries were sequenced for 25 cycles in paired-end mode on the Illumina HiSeq 2500 platform at the Fred Hutchinson Cancer Research Center Genomics Shared Resource. Paired-end reads were mapped to release r6.30 of the *D. melanogaster* genome obtained from FlyBase using Bowtie2, and to the yeast genome (SacCer3) for spike-in normalization, normalizing fly profiles to the number of yeast reads. Track screenshots were produced using the UCSC Genome browser (http://genome.ucsc.edu) [49].

#### Data accession codes

Sequencing data are available in GEO at National Center for Biotechnology Information (GSE121028).

## Acknowledgements

We thank Steven Henikoff, Welcome Bender, Jorja Henikoff, Tayler Hentges, Kristen Powers, and Isaac Brock. This work was funded by NIH grant R01GM108699 (K. Ahmad).

## Supplementary Information

**Supplementary Table 1.** New mutations of the the *vestigial* gene.

| *mutation* | *position from TSS<sup>1</sup>* | *method of induction* | *structure* |
| --- | --- | --- | --- |
| 13A | 32 – 423 | *P* mobilization | deletion and insertion of TCAT |
| CL1 | 303 – 667 | CRISPR | deletion |
| CZ2 | 263 – 344 | CRISPR, IScel cleavage | deletion |
| CL2C | 296 | CRISPR | C -> T substitution |
| DJ1 | 303 – 667 | CRISPR | deletion |
| R5 | 19614 – 20640 | CRISPR, IScel cleavage | deletion |
| R22 | 19614 – 20640 | CRISPR, IScel cleavage | deletion |

<sup>1</sup>The *vg* Transcriptional Start Site (TSS) is located at 2R:12,884,200 dm6.

**Supplementary Table 2.** Complementation^a^ of new *vestigial* alleles.

| *Df(2R)*<br>*vgD* | *Df(2R)*<br>*Exel8056* | *vg<sup>13A</sup>* | *vg<sup>CL1</sup>* | *vg<sup>CZ</sup>* | *vg<sup>CL2C</sup>* | *vg<sup>nw</sup>* | *vg<sup>R5</sup>* | *vg<sup>DJ1R22</sup>* |  |
| --- | --- | --- | --- | --- | --- | --- | --- | --- | --- |
| lethal | lethal | 6 | 5 | 1 | 4 | lethal | 0 | 5 | *Df(2R)vgD* |
|  | lethal | 5 | 5 | 1 | 3 | 6 | 0 | 6 | *Df(2R)Exel8056* |
|  |  | 5 | 5 | 1 | 0 | 5 | 0 | 5 | *vg<sup>13A</sup>* |
|  |  |  | 5 | 0 | 0 | 5 | 0 | 5 | *vg<sup>CL1</sup>* |
|  |  |  |  | 0 | 0 | 2 | 0 | 0 | *vg<sup>CZ</sup>* |
|  |  |  |  |  | 0-1 | 3 | 0 | 1 | *vg<sup>CL2C</sup>* |
|  |  |  |  |  |  | lethal | 0 | 5 | *vg<sup>nw</sup>* |
|  |  |  |  |  |  |  | 0 | 0 | *vg<sup>R5</sup>* |
|  |  |  |  |  |  |  |  | 5 | *vg<sup>DJ1R22</sup>* |

<sup>a</sup> Allelic combinations were scored in six categories based on adult wing area and wing margin scalloping from full wildtype wings (0) to complete elimination (6) of the wing, or as lethal combinations.

**Supplementary Figure 1. Native ChIP mapping of chromatin factors at the *vestigial PRE*s in Drosophila S2 cells.** Correspondence of CUT&RUN mapping of *PRE* features to bound transcription factors detected by ORGANIC native-ChIP (native-ChIP profiling was reported in [13]).

**Supplementary Figure 2. Two-step deletion of regulatory elements.** The scheme is designed to eliminate a genomic segment (white box) through a series of targeted recombination events. In (1), the YFP donor plasmid with an internally-deleted homology region is co-injeced with a CRISPR gRNA plasmid into embryos expressing Cas9. Cleavage of the chromosome by Cas9 stimulates recombination (blue lines) between the donor plasmid and the chromosome. (2) Flies with the resulting tandem duplication in the chromosome are recovered by crossing injected animals to a GAL4 driver line and screening progeny for YFP-expressing progeny. In (3), flies with the tandem duplication are crossed to animals with an heat-shock-inducible *ISce-I* gene, and the progeny are heat-shocked. These animals are crossed to a GAL4 driver line, and progeny that do not express YFP are recovered as potential intra-chromosomal recombination events that have reduced the tandem duplication (4). Deletion of the regulatory element is confirmed by PCR amplification of the homology region and sequencing.

